# Geometric coordination of endothelial invasion and vascular patterning during epithelial morphogenesis

**DOI:** 10.64898/2026.10.13.758991

**Authors:** Tsuyoshi Hirashima, Dinh Thach Lam Nguyen, Yuan Zhe Lee

## Abstract

As organs grow and remodel in three dimensions, developing vascular networks extend through the tissue while maintaining spatial coordination and establishing connections between distant sprouts. How local endothelial behavior generates vascular patterns within such deforming environments remains poorly understood. Here, using the developing mouse epididymis, we combine two-photon live microscopy, pharmacological perturbations, and mathematical modeling to investigate angiogenesis alongside epithelial tubule folding. Endothelial cells invade the organ interior while maintaining proximity to the deforming tubule. Live imaging reveals dynamic protrusions directed toward the epithelium and a characteristic sequence in which endothelial fronts approach the tubule, stop, and maintain their separation despite continued folding. ERK activity promotes endothelial extension and declines during the transition to distance maintenance. From these observations, we derive a minimal follow-and-stop rule that captures endothelial positioning relative to the changing epithelium. Extending this model to three dimensions reveals how epithelial folding can bring otherwise distant endothelial fronts together, creating opportunities for vascular connection. Explants exhibiting greater three-dimensional folding also show a higher observed frequency of endothelial connections. Our findings support a geometric framework in which local endothelial dynamics and tissue-scale deformation can jointly contribute to vascular patterning during organ morphogenesis.

## INTRODUCTION

Angiogenesis expands pre-existing vascular networks through coordinated endothelial sprouting, anastomosis, and remodeling, allowing the vasculature to develop alongside growing organs and adapt to their architecture and function^1–3^. This process requires more than an intrinsic branching program: endothelial cells (ECs) must continuously respond to growth factors, neighboring cells, extracellular-matrix cues, and mechanical signals within their surrounding tissue^4,5^. A fundamental question is how vascular growth remains spatially coordinated with organ morphogenesis when the surrounding tissue itself is continuously changing shape.

At the leading edge of angiogenic sprouts, endothelial tip cells extend dynamic filopodial protrusions that explore their local environment. VEGF-A signaling through VEGFR2 promotes endothelial migration and sprout extension, whereas DLL4–Notch signaling regulates tip-cell selection among neighboring ECs^6–9^. However, tip and stalk identities are not fixed. Individual ECs can exchange positions and compete for the leading position^10^, while live imaging and mathematical modeling have revealed extensive cell rearrangement, overtaking, and coordinated movement within growing sprouts^11–13^. Angiogenic sprouting is therefore a dynamic and adaptive process in which individual cell behaviors collectively determine vascular growth.

Beyond molecular signaling, angiogenic sprouts must navigate the physical architecture of their surrounding tissue. Mechanical properties, including matrix stiffness, tissue tension, and fluid-derived forces, influence endothelial migration and vascular morphogenesis^14–17^. Structural features, such as matrix organization and surface topography, further influence vascular organization^18–20^. Together with attractive and repulsive biochemical cues^21–24^, these physical factors shape the paths available to growing vessels. Vascular patterning thus emerges from the interplay between biochemical signaling, tissue mechanics, and geometry.

An additional challenge arises during three-dimensional organogenesis, when the architecture of the surrounding tissue is itself changing. Much of our mechanistic understanding of angiogenic guidance comes from experimentally tractable systems, including the postnatal mouse retina and zebrafish vasculature^6,7,10,25^. During the development of complex organs, however, neighboring epithelial structures elongate, bend, and fold, continuously altering the spatial relationships between extending endothelial fronts and their surroundings. These geometric changes may also bring previously distant vascular fronts into proximity, creating new opportunities for connection. Yet whether a morphogenetic epithelial structure can serve as a dynamic positional reference for ECs, and how its changing geometry contributes to vascular connectivity, remain poorly understood.

Here, we investigate how local endothelial behaviors give rise to vascular patterns during three-dimensional organ morphogenesis. We use the developing mouse epididymis, in which the Wolffian duct-derived epithelial tubule undergoes extensive elongation and folding as endothelial cells invade the surrounding interstitium^26–29^. This setting therefore allows vascular extension and epithelial deformation to be followed and quantified within the same intact organ. Combining whole-mount imaging, ex vivo two-photon live imaging, pharmacological perturbations, and computational modeling, we show that endothelial fronts remain spatially associated with the deforming tubule and repeatedly extend and retract protrusions toward it. As the tubule folds, endothelial fronts exhibit a simple follow-and-stop behavior: they approach the epithelium and subsequently maintain a characteristic separation, accompanied by a decline in ERK activity. A minimal geometric model demonstrates how this local behavior can maintain endothelial proximity to the changing tubule, while three-dimensional epithelial folding brings otherwise separated endothelial fronts into positions favorable for connection. Here, “geometric” refers to the spatial relationship between endothelial fronts and the changing epithelial architecture and does not presuppose the molecular or physical mechanism by which this relationship is sensed. Stochastic changes in endothelial migration direction provide an alternative route to such encounters. Consistent with these predictions, explants exhibiting greater three-dimensional folding also show a higher observed frequency of endothelial connections. Together, our findings suggest that vascular network organization can emerge from the interplay between local endothelial dynamics and tissue-scale epithelial deformation, providing a simple geometric framework for understanding angiogenesis during organ morphogenesis.

## RESULTS

### Vascular invasion accompanies epididymal tubule folding while preserving EC–tubule proximity

We first examined the spatial relationship between ECs and the epithelial tubule during epididymal morphogenesis in developing murine embryos. Whole-mount immunofluorescence for CD31 and PAX2 was performed on intact epididymides at E15.5, E16.5, and E17.5, spanning the period in which the initially straighter epididymal tubule develops progressively deeper folds^26–28^. In the peripheral region adjacent to the organ surface, a connected CD31-positive vascular plexus was already evident at E15.5 and remained broadly similar at later stages. In contrast, the vascular pattern in the deep interior of the epididymis changed markedly. At E15.5, ECs were concentrated beneath the organ surface; by E16.5, endothelial extensions entered clefts associated with the folding tubule; and by E17.5, EC networks extended further into the interior and formed connections across the folded epithelial geometry (**Figure 1A; Video 1**). Thus, vascular invasion of the deep epididymis occurs concurrently with epithelial tubule morphogenesis.

**Figure 1.**
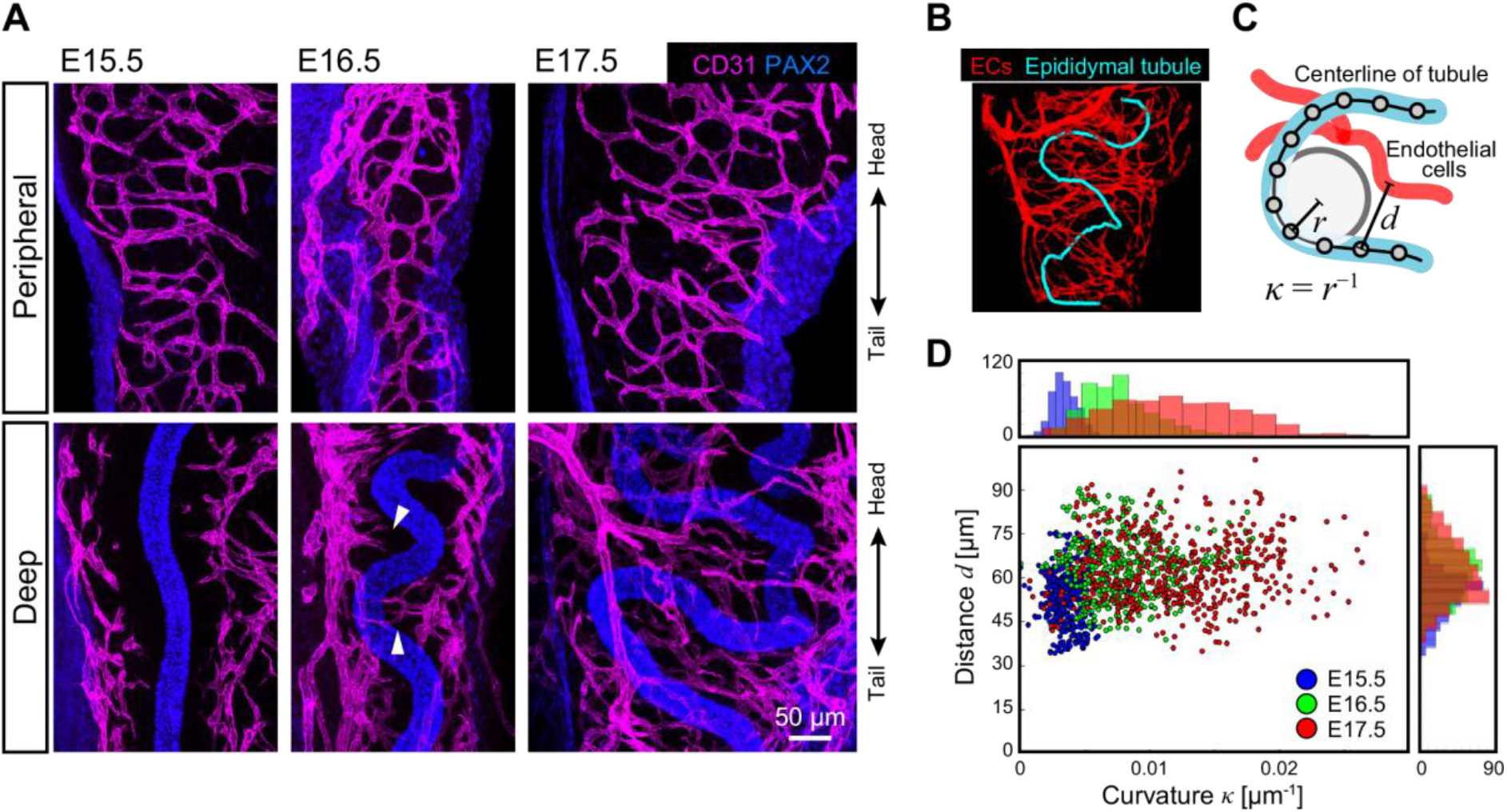
Spatial coordination of endothelial cells and the epithelial tubule during epididymal development. (A) Maximum-intensity projections of whole-mount immunofluorescence for CD31 (magenta) and PAX2 (blue) at E15.5, E16.5, and E17.5. Upper panels show peripheral regions; lower panels show deeper regions. Arrowheads indicate endothelial extensions entering the interior adjacent to tubule folds. Scale bar, 50 µm. (B) Representative three-dimensional segmentation of the EC network (red) and epididymal tubule/centerline (cyan). (C) Schematic defining local curvature *κ*=1/*r*, where *r* is the local radius of curvature of the tubule centerline, and *d*, the minimum distance from a centerline position to the EC network. (D) Scatter plots of local tubule curvature and EC-tubule distance at the indicated embryonic stages, with marginal histograms. n>420 centerline positions pooled from N=5 embryos. Pearson’s correlation is −0.018, −0.038, and 0.042 at E15.5, E16.5, and E17.5, respectively.

To quantify this spatial coordination in three dimensions, we segmented the CD31-positive vasculature and PAX2-positive tubule and extracted the epithelial centerline (**Figure 1B; Video 2**). For each centerline position, we measured the local tubule curvature, *κ*, defined as the inverse local radius of curvature, and the minimum Euclidean distance, *d*, to the segmented EC network (**Figure 1C**). As expected, the curvature distribution broadened and shifted toward higher values from E15.5 to E17.5, reflecting progressive tubule folding (**Figure 1D**). In contrast, the distribution of EC–tubule distance remained similar from E15.5 to E17.5 despite the progressive increase in tubule curvature, and showed essentially no correlation with local curvature. These data indicate that, despite substantial changes in epithelial geometry, the developing vascular network remains spatially registered to the tubule rather than simply occupying fixed positions within the organ.

### Vascular depletion does not prevent epididymal tubule folding

We next examined whether perturbation of ECs affects epididymal tubule folding. Because VEGF-A/VEGFR2 signaling is a major regulator of endothelial survival, migration, and angiogenic sprouting^2,6^, we first examined the spatial distribution of VEGF-A and VEGFR2 in E16.5 epididymides. VEGF-A immunoreactivity was strongest in the epithelial tubule but was also detected throughout the surrounding interstitium, whereas VEGFR2 was enriched in CD31-positive ECs and was not detectably enriched in the epithelial tubule (**Figures 2A,B**). This spatial distribution is consistent with the established role of VEGFR2 as a principal endothelial receptor mediating VEGF-A-dependent angiogenic responses^6,30^.

**Figure 2.**
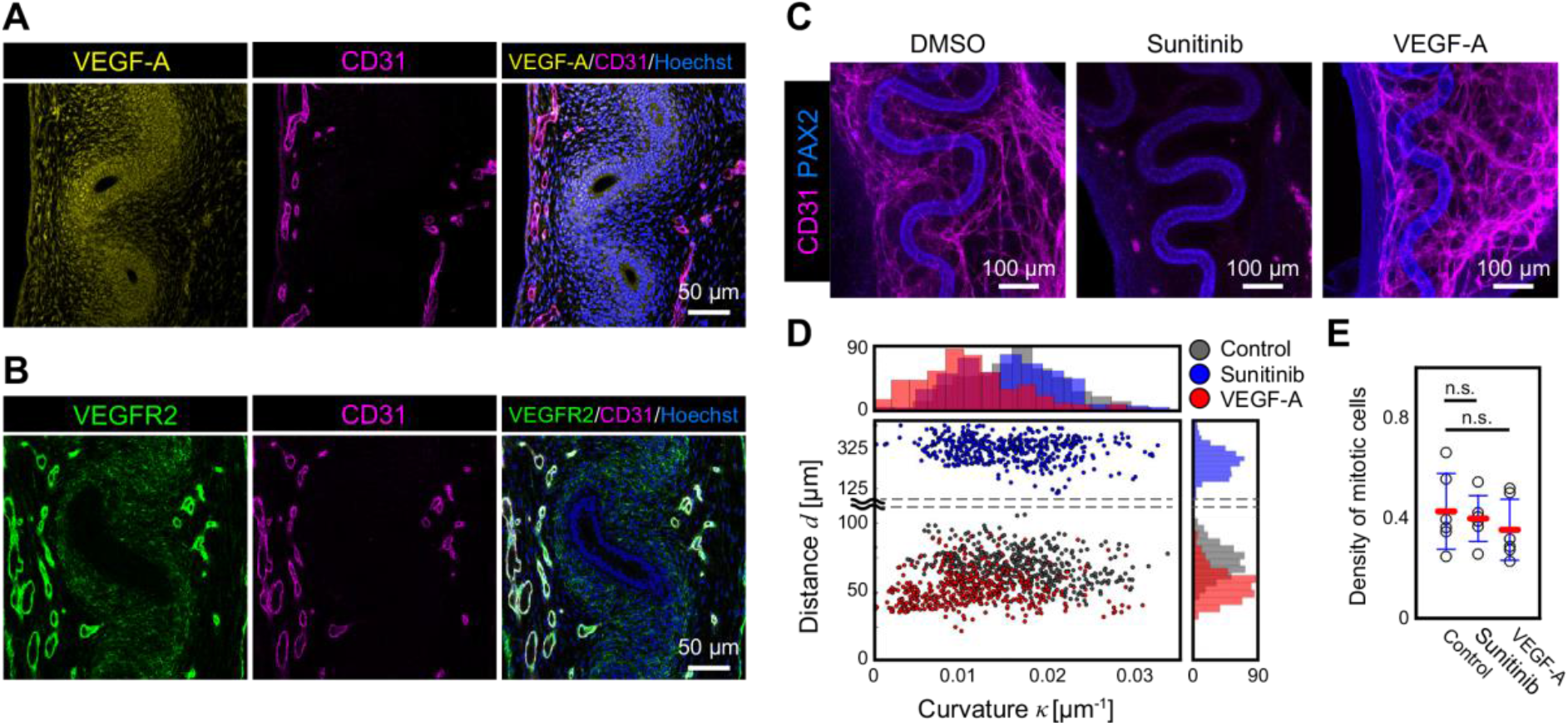
Effects of angiogenic perturbation on vascular patterning and epididymal tubule morphology. (A,B) Whole-mount immunofluorescence of E16.5 epididymides for VEGF-A (yellow; A) or VEGFR2 (green; B), together with CD31 (magenta) and Hoechst nuclear staining (blue). Scale bars, 50 µm. (C) Maximum-intensity projections of CD31 (magenta) and PAX2 (blue) in E16.5 epididymides cultured ex vivo for 24 h with DMSO vehicle, sunitinib (5 µM), or recombinant VEGF-A (100 ng/mL). Scale bars, 100 µm. (D) Scatter plots of local tubule curvature, *κ*, and EC–tubule distance, *d*, under the indicated conditions. Marginal histograms show the distributions of *κ* and *d*. n>460 centerline positions pooled from N=6 embryos. (E) Density of phospho-histone H3-positive mitotic cells within the epididymal tubule under the indicated conditions. Points represent individual embryos; bars indicate mean ± s.d.; N=6 embryos per condition. One-way ANOVA followed by Dunnett’s multiple-comparison test versus vehicle control; sunitinib, p=0.88; VEGF-A, p=0.51.

We then cultured E16.5 epididymides ex vivo for 24 h with vehicle, the multi-target receptor tyrosine kinase inhibitor sunitinib^31,32^, or recombinant VEGF-A. Sunitinib markedly reduced the CD31-positive vascular network, leaving sparse EC clusters predominantly near the organ periphery, whereas VEGF-A treatment increased the abundance of CD31-positive vascular structures throughout the tissue (**Figure 2C**). Consistent with these changes in EC distribution, the EC–tubule distance, *d*, increased following sunitinib treatment and decreased following VEGF-A treatment (**Figure 2D**).

Despite the marked reduction in ECs following sunitinib treatment, the distribution of tubule curvature remained similar to that of the vehicle control (**Figure 2D**). VEGF-A treatment shifted the curvature distribution toward lower values, although tubule folding was still evident (**Figures 2C,D**). Because epithelial proliferation contributes to epididymal tubule elongation and folding^26,27^, we next quantified phospho-histone H3-positive mitotic cells within the tubule. Mitotic cell density was not detectably altered by either sunitinib or VEGF-A treatment (**Figure 2E**). Together, these results show that substantial depletion of the developing vasculature does not prevent epididymal tubule folding over this developmental interval.

### Endothelial filopodia dynamically probe the epididymal tubule

We next examined the morphology and dynamics of endothelial fronts relative to the epithelial tubule. To visualize EC membranes and the tubule simultaneously, we used Hoxb7-myr-Venus transgenic mice, in which membrane-anchored Venus is expressed under the control of the Hoxb7 promoter^33^. In the developing epididymis, Venus fluorescence also outlined CD31-positive vascular structures, enabling simultaneous visualization of EC membranes and the epithelial tubule (**Figure 3A**).

**Figure 3.**
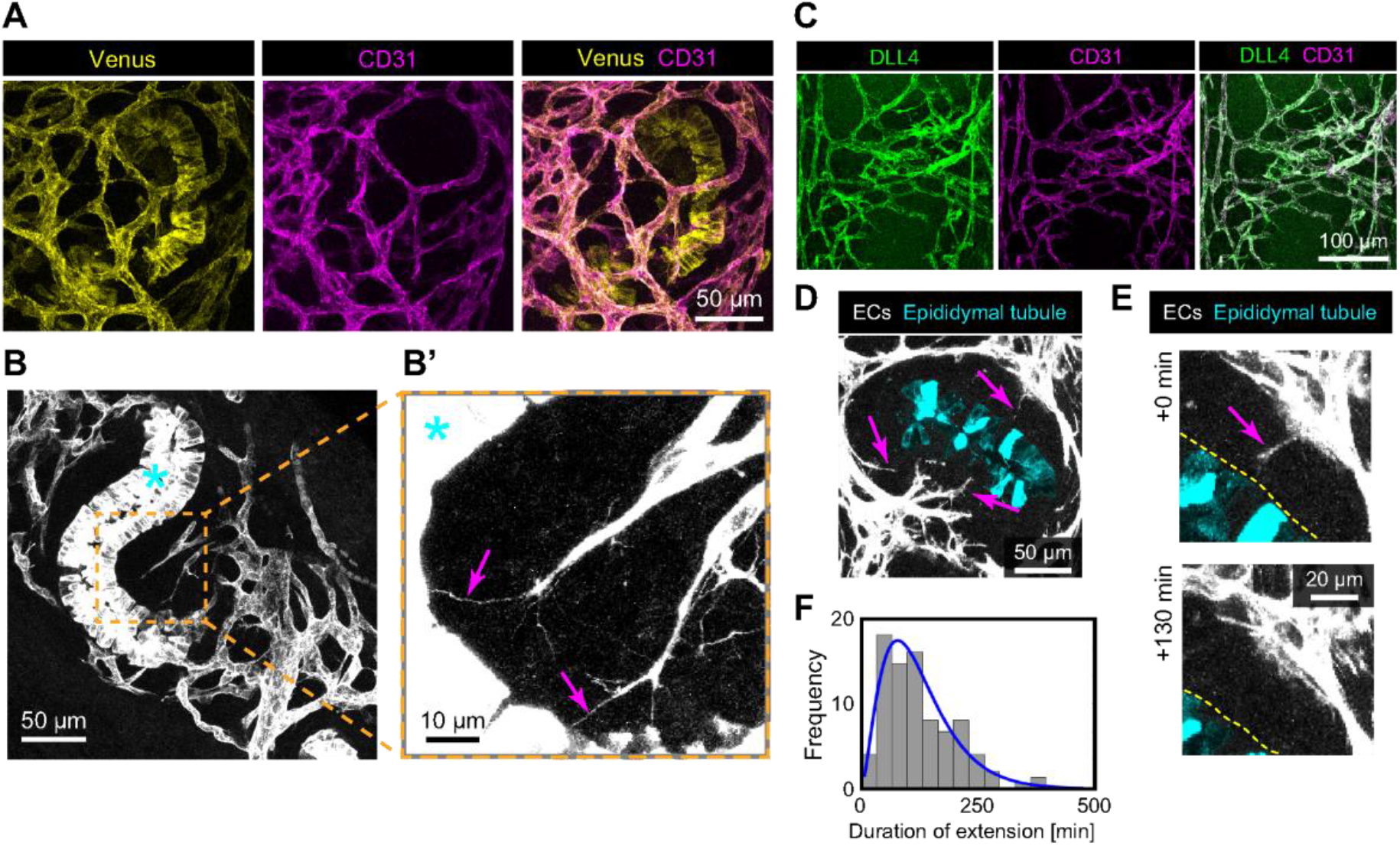
Dynamic endothelial protrusions extend toward the epididymal tubule. (A) Maximum-intensity projections of Venus fluorescence (yellow) and CD31 immunofluorescence (magenta) in an E16.5 Hoxb7-myr-Venus epididymis. Venus fluorescence labels both the epithelial tubule and endothelial membranes. Scale bar, 50 µm. (B,B’) Contrast-enhanced Hoxb7-myr-Venus images of an E17.5 epididymis. The boxed region in (B) is enlarged in (B’). Asterisks indicate the epithelial tubule, and arrows indicate thin endothelial protrusions extending toward the tubule. Scale bars, 50 µm (B) and 10 µm (B’). (C) Maximum-intensity projections of DLL4 (green) and CD31 (magenta) immunofluorescence in an E17.5 epididymis. Scale bar, 100 µm. (D,E) Representative snapshots from ex vivo two-photon time-lapse imaging of Hoxb7-myr-Venus epididymides. The epididymal tubule is pseudocolored cyan, and magenta arrows indicate endothelial protrusions. (E) Representative example of protrusion extension followed by retraction. Scale bars, 50 µm (D) and 20 µm (E). (F) Distribution of endothelial protrusion-extension durations. The blue curve indicates a gamma distribution fitted to the measured durations by maximum likelihood. The fitted shape and scale parameters were *k*=2.86 and *θ*=43.8 min, respectively. n=41 extension events from N=3 embryos.

Two-photon imaging of optically cleared E17.5 epididymides revealed thin membrane protrusions extending from endothelial fronts toward the epithelial tubule (**Figures 3B,B’**). Following local contrast enhancement, individual protrusions could be traced for tens of micrometers, with their distal tips closely apposed to the tubule surface. Based on their slender morphology and extension from endothelial fronts, these structures resembled endothelial filopodia, which are known to mediate environmental sensing and directional migration during angiogenic sprouting^6,34^.

Because these protrusions resembled those associated with angiogenic tip-cell behavior, we next examined the distribution of DLL4, a key regulator of tip-cell selection through Notch signaling^7,8^. DLL4 immunoreactivity was broadly detected across the CD31-positive angiogenic network in the developing epididymis (**Figure 3C**), rather than being restricted to a small subset of morphologically distinct leading cells.

To determine whether these tip-associated protrusions were stable or dynamically remodeled, we performed two-photon time-lapse imaging of Hoxb7-myr-Venus epididymides. Filopodia-like endothelial protrusions repeatedly extended toward and retracted from the epithelial tubule (**Figures 3D,E; Video 3**). The durations of individual extension phases were heterogeneous and showed a broad right-skewed distribution with a peak away from zero. A gamma fit to the extension durations yielded a shape parameter *k* of approximately 2.9 and a characteristic scale *θ* of approximately 44 min (**Figure 3F**). Notably, *k*>1 distinguishes this behavior from a simple memoryless termination process and indicates temporally structured stochastic dynamics^35,36^, in which the propensity to terminate extension increases with elapsed time. Thus, endothelial fronts undergo repeated cycles of protrusion extension and retraction toward the tubule, indicating a dynamic rather than persistent interaction with the local epithelial environment.

### Endothelial fronts follow tubule bending and stop at a characteristic separation

The spatial proximity between ECs and the epithelial tubule, together with the repeated extension of endothelial protrusions toward the tubule, raised the question of how this relationship is maintained as the tubule continuously changes shape. Long-term ex vivo imaging was therefore used to follow EC position and epithelial tubule geometry during active folding (**Figure 4A; Video 4**). In E15.5 epididymides, the epithelial fold progressively deepened over the imaging period, while a neighboring endothelial front advanced toward the same region. To separate tubule deformation from EC–tubule positioning, we defined two geometric quantities (**Figure 4B**): *H*, the displacement of the target tubule apex relative to the line connecting the opposing fold apices, and *D*, the distance between the EC tip and the target tubule apex.

**Figure 4.**
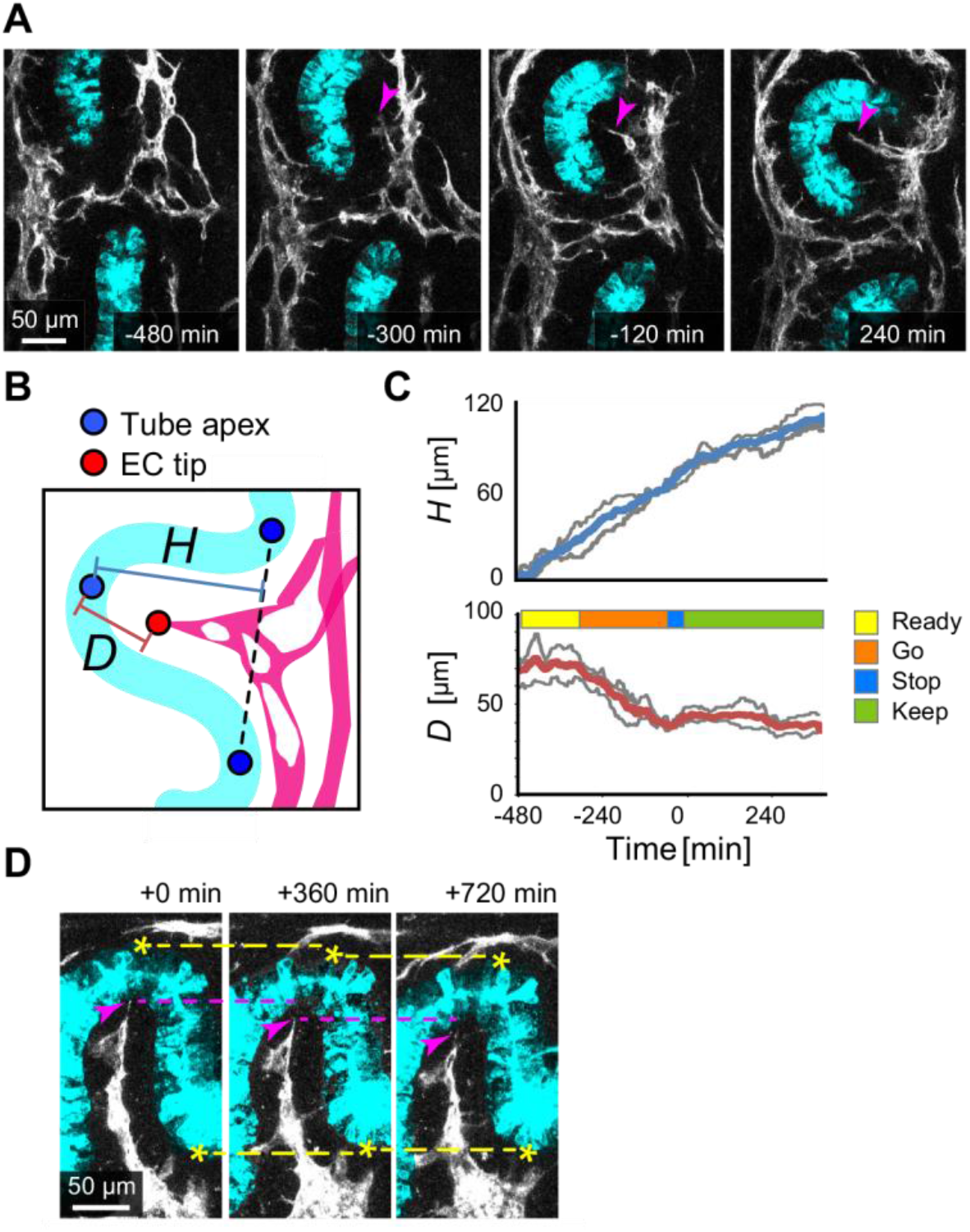
Endothelial fronts approach the folding tubule and stabilize their separation. (A) Representative time-lapse sequence of an E15.5 epididymis showing an endothelial front advancing toward the cleft of a folding epithelial tubule. The epithelial tubule is pseudocolored cyan, and arrowheads indicate the endothelial front. Times are relative to the onset of Stop phase. Scale bar, 50 µm. (B) Schematic defining the tubule apex displacement, *H*, and the EC-tip-to-tubule-apex distance, *D*. Blue points indicate tubule apices, and the red point indicates the endothelial tip. (C) Time courses of *H* (top) and *D* (bottom). Gray lines indicate individual samples, and colored lines indicate the mean (N=3 embryos). The colored bar indicates the operationally defined Ready, Go, Stop, and Keep phases. (D) Representative example of a local change in tubule geometry during ex vivo imaging accompanied by retraction of the nearby endothelial front. Yellow dashed lines indicate fixed spatial references, and arrowheads indicate the endothelial front. Scale bar, 50 µm.

*H* increased progressively throughout the observation period, consistent with continued tubule folding (**Figure 4C, top**). In contrast, *D* showed a distinct temporal progression (**Figure 4C, bottom)**. During an initial ‘Ready’ phase, *D* remained relatively stable despite the continued increase in *H*. This was followed by a ‘Go’ phase, during which the EC front approached the tubule and *D* decreased substantially. Once *D* reached approximately 30–40 µm in these recordings, further approach slowed and the separation remained within a similar range despite continued changes in tubule geometry. We operationally termed this transition the ‘Stop’ phase, followed by a ‘Keep’ phase in which the reduced EC–tubule separation was maintained. Thus, tubule deformation and EC extension were not coupled in a simple one-to-one manner; instead, the EC front exhibited a delayed approach followed by stabilization of its position relative to the tubule.

We also observed occasional local rearrangements during prolonged ex vivo imaging, in which deformation of a nearby epithelial fold was accompanied by retraction of the associated endothelial front (**Figure 4D; Video 5**). In these cases, the EC tip shifted back as the tubule geometry changed, rather than maintaining its previous position. Although such events were infrequent and were not quantified at the population level, they were consistent with the idea that EC–tubule separation is dynamically maintained as the tubule deforms.

Together with the repeated extension and retraction of endothelial protrusions toward the tubule, these observations suggest that endothelial fronts dynamically adjust their position relative to the changing epithelial geometry. This behavior can be summarized by a minimal “follow-and-stop” rule: EC fronts advance when sufficiently separated from the tubule and reduce their forward extension after entering a local proximity range.

### ERK activity promotes endothelial extension and declines near the epithelial tubule

The transition from active approach to stable EC–tubule separation suggested that endothelial motility is actively modulated as EC fronts approach the epithelial tubule. ERK is a central downstream effector of growth-factor signaling and exhibits dynamic activity during endothelial migration and angiogenesis^37,38^. We therefore examined whether altering ERK signaling affects endothelial extension in the developing epididymis. Treatment with the MEK inhibitor PD0325901 markedly reduced EC extension into the deep region of the epididymis and diminished the bridging vascular patterns observed in vehicle-treated samples (**Figure 5A**). In contrast, EGF was used as an upstream activator of the ERK pathway, and its stimulation increased the abundance of endothelial extensions. These opposing responses suggested a positive relationship between ERK pathway activity and endothelial extension.

**Figure 5.**
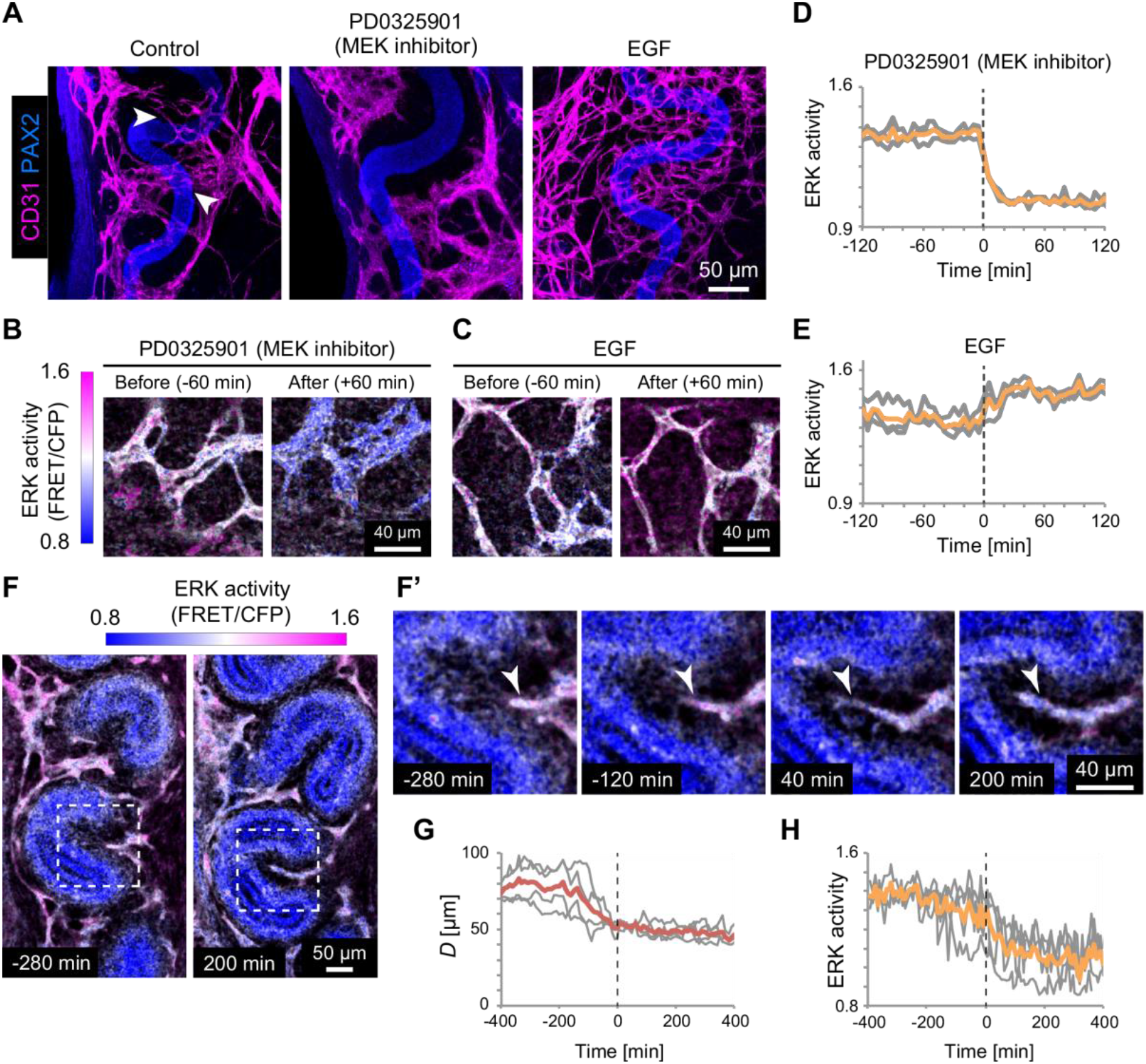
ERK activity promotes endothelial extension and declines during distance maintenance. (A) Maximum-intensity projections of CD31 (magenta) and PAX2 (blue) after 12 h treatment with vehicle, the MEK inhibitor PD0325901 (100 nM), or EGF (10 ng/mL). Arrowheads in the vehicle control indicate endothelial extensions bridging around or across the folded tubule. Scale bar, 50 µm. (B,C) Representative ERK activity maps before and after treatment with PD0325901 (B) or EGF (C). Scale bars, 40 µm. (D, E) Time courses of ERK activity following PD0325901 (D) or EGF (E) treatment. Gray lines indicate individual samples, and orange lines indicate the mean. Time zero indicates the time of treatment. N=3 embryos. (F,F’) Representative ERK activity maps during endothelial approach toward the epithelial tubule. (F’) Enlarged view of the boxed region in (F). Time zero is aligned to the onset of the Stop phase. Scale bars, 50 µm (F) and 40 µm (F’). (G,H) Time courses of EC-tip-to-tubule distance, *D* (G), and ERK activity (H), aligned to the onset of the Stop phase. Gray lines indicate individual embryos, and colored lines indicate the mean. N=4 embryos.

To directly confirm the effects of these treatments on ERK signaling in living tissue, we used transgenic ERK activity reporter mice, in which changes in ERK kinase activity can be monitored as a ratiometric fluorescence signal^39^. Two-photon imaging revealed a rapid decrease in ERK activity following PD0325901 treatment, whereas EGF stimulation produced a clear increase (**Figures 5B–E**). Thus, the opposing effects on endothelial morphology were accompanied by corresponding decreases and increases in ERK activity.

Having established that ERK activity could be monitored dynamically in the explant, we then examined how ERK signaling changed as EC fronts approached the tubule and entered the stop phase. Live ERK activity maps revealed spatially heterogeneous signaling within the developing vascular network (**Figure 5F; Video 6**). To compare this transition across samples, time-lapse sequences were aligned to the onset of the Stop phase, defined as *T*=0 from the EC–tubule distance trajectory. As expected from the previous results (**Figure 4**), *D* decreased during the approach period and subsequently stabilized (**Figures 5F’,G**). ERK activity was relatively high while the EC front was approaching the tubule but decreased around the transition to the Stop phase, remaining lower as *D* stabilized (**Figure 5H; Video 7**). The decrease in ERK activity therefore coincided temporally with the transition from active endothelial extension to maintenance of EC–tubule separation.

Together, the perturbation and live-imaging experiments identify ERK activity as a dynamic regulator of endothelial motility. Pharmacological suppression of ERK signaling inhibits EC extension, whereas ERK activation enhances endothelial outgrowth. Conversely, endogenous ERK activity decreases as EC fronts transition from approach to distance maintenance. Thus, ERK functions as a tunable motility controller, with higher activity favoring endothelial advance and lower activity accompanying braking and stabilization of the EC front.

### A geometric model links endothelial follow-and-stop dynamics and epithelial folding to shape vascular connectivity

Our live-imaging experiments showed that endothelial fronts approach the folding epithelial tubule and subsequently maintain a characteristic separation, accompanied by a decline in ERK activity. We next asked whether this local behavior could explain not only the spatial relationship between ECs and the tubule but also the formation of vascular connections within the developing organ. To address this question, we developed a minimal model in which EC fronts extend toward a deforming epithelial tubule according to a local follow-and-stop rule (**Figure 6A**). Each front advances at an intrinsic speed, v_tip_, and senses the tubule within a forward cone defined by its half-angle, α, representing the spatial range of filopodial exploration (**Figure 6A’**). Advancement stops when the tubule enters a prescribed proximity range, allowing the model to represent endothelial approach and distance maintenance without specifying the underlying molecular sensing mechanism.

**Figure 6.**
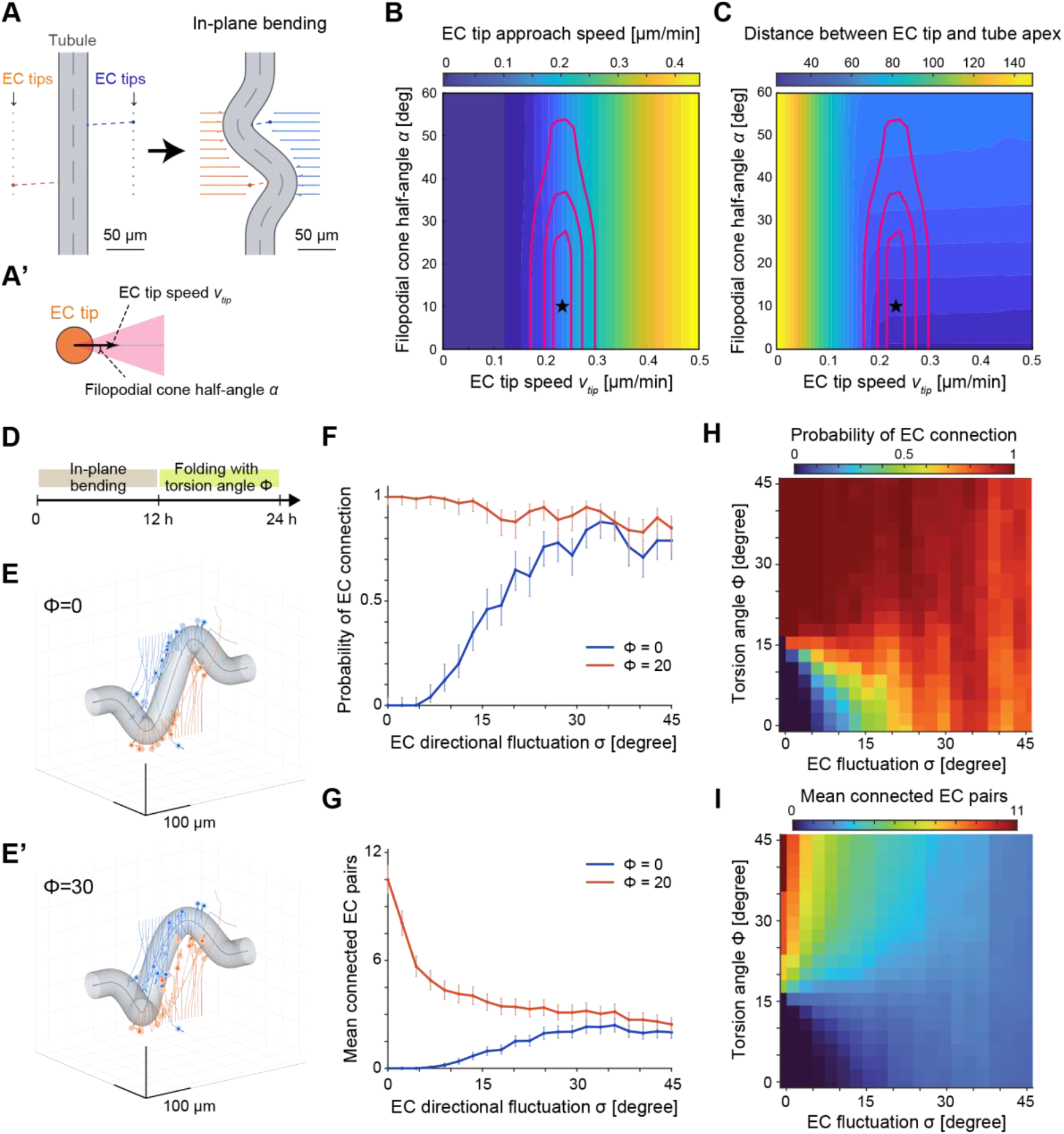
Computational analysis of endothelial approach and connection during epithelial tubule folding. (A,A’) Schematic of the planar bending model, showing EC fronts initially positioned on opposite sides of the epithelial tubule (A) and definition of EC tip extension speed v_tip_ and filopodial sensing-cone half-angle α. Scale bars, 50 µm. (B, C) Parameter maps of EC tip approach speed (B) and late EC tip–tubule apex distance (C) as functions of v_tip_ and α. Magenta contours indicate the optimization criteria based on experimentally measured approach speed and maintained EC–tubule distance, respectively. The asterisk indicates the reference parameter combination selected by minimizing the combined mismatch between simulations and experimental measurements. (D) Prescribed morphogenetic schedule comprising 12 h of in-plane bending followed by 12 h of folding controlled by the torsion parameter Φ. (E,E′) Representative three-dimensional configurations of the epithelial tubule and EC fronts at 24 h for Φ=0 (E) and Φ=30 (E′). Scale bars, 100 µm. (F,G) Probability of at least one connection between opposing EC fronts (F) and mean number of connected EC pairs (G) at 24 h as functions of EC directional fluctuation σ, for Φ=0 and Φ=20. Error bars indicate 95% Wilson confidence intervals (F) and mean ± SEM (G), based on 100 independent stochastic simulations per condition. (H,I) Parameter maps of connection probability (H) and mean number of connected EC pairs (I) at 24 h as functions of Φ and σ. Each parameter combination was evaluated using 100 stochastic realizations.

We first simulated planar tubule bending and varied v_tip_ and α to identify parameter combinations consistent with our experimental observations (**Figures 6B,C**). Comparing simulated approach speed and late EC–tubule separation with the experimental measurements (**Figure 4C**) yielded a reference combination of v_tip_=0.24 µm/min and α=10°. We then extended the model further, following the previously described progression from in-plane bending to more complex epididymal tubule folding^27^. The simulated tubule underwent 12 h of in-plane bending followed by 12 h of additional out-of-plane folding, controlled by the parameter Φ (**Figure 6D**). We also introduced stochastic changes in EC migration direction, with their timing determined by the experimentally measured protrusion-extension durations (**Figure 3F**) and their angular variability controlled by σ.

These two model parameters represent distinct routes to endothelial connection: epithelial folding can reposition opposing fronts, whereas directional fluctuations allow ECs to explore alternative paths. To examine how these effects compete, we developed a simplified analytical approximation. For a given connection probability *p*, their relationship can be expressed as

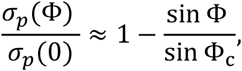

where *σ_p_*(0) is the directional fluctuation required without out-of-plane folding, and Φ_c_ is the geometric connection threshold. This relation predicts that epithelial folding and directional fluctuations can compensate for one another: greater folding reduces the stochastic exploration required for EC connection, potentially eliminating this requirement near Φ_c_. This approximation applies to the low-fluctuation transition boundary, rather than the entire parameter space (**Supplementary Note**).

We next examined this competition through stochastic simulations of EC extension and connection over 24 h (**Figures 6E,E’; Video 8**). In the planar case (Φ=0), connections were rare without directional fluctuations, but their probability increased substantially with σ (**Figure 6F**). In contrast, out-of-plane folding (Φ=20) produced a high connection probability even with little directional fluctuation. A different pattern emerged when we counted the number of connected EC pairs (**Figure 6G**). With out-of-plane folding, multiple opposing fronts connected at low σ, but their number decreased markedly as directional fluctuations increased. Under planar bending, increasing σ instead enabled connections that were largely absent without stochastic exploration. Thus, directional fluctuations can facilitate an initial encounter while simultaneously limiting the formation of multiple connections when epithelial geometry already favors their formation.

To determine whether these behaviors extended across the parameter space, we systematically varied Φ and σ (**Figures 6H,I**). The connection-probability map revealed a trade-off between folding and directional fluctuations: increasing either parameter could promote at least one connection when the other was limited (**Figure 6H**), consistent with the trend predicted by the reduced analytical approximation (**Supplementary Note**). However, the map of connected-pair numbers revealed a more restricted region of high connectivity, characterized by substantial out-of-plane folding and relatively small directional fluctuations (**Figure 6I**). Together, these results distinguish the conditions required for a single endothelial encounter from those favoring multiple connections.

Our model therefore links local endothelial behavior to vascular connectivity at the tissue scale. The follow-and-stop rule maintains EC proximity to the deforming epithelial tubule, while three-dimensional folding repositions opposing fronts to create opportunities for connection. Directional fluctuations provide an alternative means of encountering otherwise separated fronts; however, extensive stochastic exploration is not required when epithelial geometry brings multiple fronts into favorable positions. These findings suggest that organ morphogenesis can contribute directly to vascular network organization by controlling the spatial opportunities for endothelial connection.

### Three-dimensional epithelial folding is associated with a higher observed frequency of endothelial connections

Our computational model suggested that out-of-plane epithelial folding can bring opposing endothelial fronts into proximity and facilitate their connection. To examine this relationship experimentally, we compared two ex vivo culture conditions: interface culture, in which epididymides were maintained at an air–liquid interface, and submerged culture, in which they were fully immersed in medium (**Figure 7A**).

**Figure 7.**
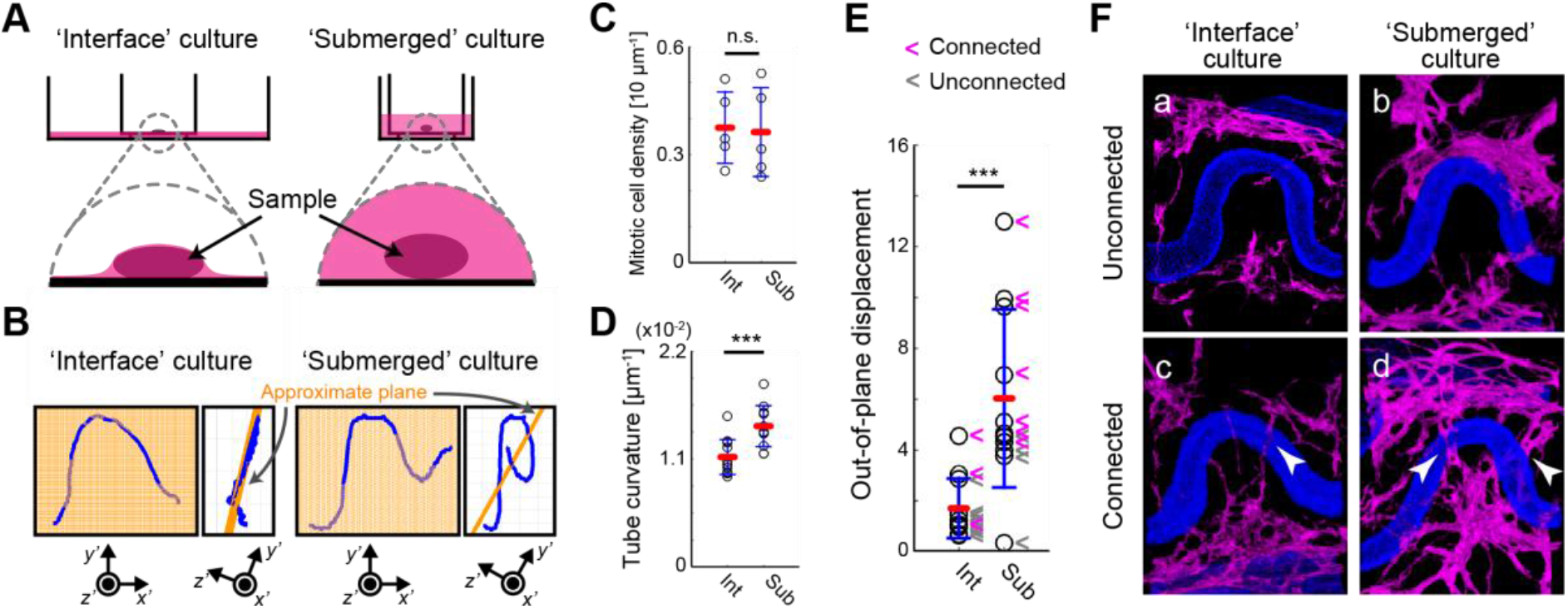
Submerged culture enhances three-dimensional epididymal folding and a higher observed frequency of endothelial connections. (A) Schematics of ex vivo interface (left) and submerged culture (right). Epididymides were maintained at an air–liquid interface or fully immersed in culture medium, respectively. (B) Representative epithelial tubule centerlines after 24 h of culture, shown in two orthogonal views. Orange lines indicate the approximate best-fit planes. (C) Density of phospho-histone H3-positive mitotic epithelial cells under the two culture conditions. N = 5 epididymides per condition. Each point in represents one epididymis; bars indicate mean ± SD. Two-sided Mann–Whitney U test; p=0.8413. (D) Mean epithelial tubule curvature after 24 h of culture. N=11 epididymides per condition. Each point in represents one epididymis; bars indicate mean ± SD. Two-sided Mann–Whitney U test; p=0.00186. (E) Mean out-of-plane displacement of the epithelial tubule centerline relative to its best-fit plane. Magenta and gray arrowheads indicate individual epididymides with connected and unconnected endothelial structures, respectively. Endothelial connections were observed in 27.3% (3/11) of interface-cultured and 63.6% (7/11) of submerged-cultured epididymides. The difference in connection frequency was not statistically significant (two-sided Fisher’s exact test, p=0.198; risk difference, 36.4 percentage points; 95% CI, −4.3 to 63.9 percentage points). Each point in represents one epididymis; bars indicate mean ± SD. Two-sided Mann–Whitney U test; p=0.00186. (F) Representative whole-mount immunofluorescence images showing unconnected (a,b) and connected (c,d) endothelial structures under interface and submerged culture conditions. PAX2-positive epithelial tubules are shown in blue, and CD31-positive endothelial cells in magenta. White arrowheads indicate endothelial connections.

After 24 h, the two conditions produced distinct epithelial morphologies (**Figure 7B**). Tubules in interface culture exhibited predominantly planar bending, consistent with constraints imposed by medium surface tension, whereas those in submerged culture developed more pronounced three-dimensional folds. Despite comparable epithelial mitotic-cell densities (**Figure 7C**), submerged culture resulted in greater mean tubule curvature (**Figure 7D**) and out-of-plane deformation (**Figure 7E**). These results established two culture conditions with distinct epithelial geometries without a detectable difference in epithelial proliferation.

We next examined whether these morphological differences were accompanied by changes in vascular connectivity. Endothelial connections were observed in 27.3% (3/11) of epididymides under interface culture and 63.6% (7/11) under submerged culture **(Figure 7E; Video 9**); this difference did not reach statistical significance (two-sided Fisher’s exact test, p=0.198). Representative images illustrate the vascular configurations surrounding epithelial folds under both conditions (**Figure 7F**). Thus, the culture condition associated with more extensive three-dimensional folding also showed a higher observed frequency of endothelial connections.

These findings are consistent with our model prediction that out-of-plane epithelial folding creates opportunities for opposing endothelial fronts to encounter one another. Although the culture conditions may influence factors beyond tissue geometry, the association between epithelial folding and endothelial connectivity supports the proposed contribution of organ morphogenesis to vascular network organization.

## DISCUSSION

In this study, we identify a simple spatial relationship that coordinates angiogenesis with epithelial morphogenesis in the murine developing epididymis. As the epithelial tubule changes from a relatively straight structure into a deeply folded three-dimensional geometry, ECs invade the tissue interior while remaining within a constrained distance of the tubule. Live imaging further resolves a dynamic sequence underlying this spatial pattern: endothelial fronts repeatedly extend protrusions toward the tubule, approach the developing fold, and then reduce their forward extension while maintaining their relative position. ERK activity promotes endothelial extension and decreases around the transition to this maintenance phase. These observations can be reduced to a simple follow-and-stop rule that links local endothelial behavior to tissue-scale vascular organization. Together, our findings support a view in which the morphogenetic epithelial tubule acts as a moving local reference for vascular invasion, rather than angiogenesis occurring within a geometrically fixed tissue environment.

This framework complements the classical view of angiogenic guidance by VEGF. In the retina, VEGF gradients provide directional information to endothelial tip-cell filopodia, while DLL4–Notch signaling regulates competition for the tip position^6,7,10^. Recent work in zebrafish further showed that neural progenitor-derived Apelin can provide local paracrine guidance for endothelial tip-cell migration and filopodial behavior while modulating ERK signaling^40^. In the developing epididymis, perturbation of VEGF signaling strongly alters vascular abundance and penetration, and modulation of the ERK pathway changes endothelial extension. However, VEGF-A immunoreactivity is broadly distributed at the spatial resolution of our assay, whereas the EC network remains locally registered to a highly dynamic epithelial geometry. We therefore distinguish two conceptually different, but interacting, levels of regulation: growth-factor signaling provides angiogenic competence and motility, whereas the tubule-associated microenvironment may provide local positional information that determines where endothelial fronts advance and where they stabilize. The molecular or physical nature of this positional information remains to be identified.

Dynamic endothelial protrusions provide a plausible cellular interface for such local sensing. Endothelial filopodia are well established as exploratory structures that sample extracellular guidance information, while computational models have linked tip-cell selection, migration, and fusion to collective sprout dynamics^6,12,13^. In the epididymis, protrusions repeatedly extend toward and retract from the epithelial tubule rather than forming a persistent attachment. Their extension durations are also inconsistent with a simple memoryless termination process, suggesting temporally structured stochastic dynamics. Together, these observations are compatible with iterative sampling of a local environment whose configuration itself changes during morphogenesis. Importantly, our data do not require the epithelial tubule to serve directly as a physical substrate for EC migration. Endothelial protrusions could instead respond to short-range soluble signals, extracellular-matrix organization, mesenchymal deformation, mechanical cues, or combinations of these factors. The follow-and-stop concept therefore describes the geometry and dynamics of the interaction without presupposing its molecular implementation.

ERK dynamics provide a signaling correlate of the transition between extension and maintenance. ERK signaling has been implicated in dynamic endothelial behaviors during regenerative angiogenesis^37^, and our pharmacological experiments similarly show that suppressing ERK activity limits endothelial extension, whereas pathway activation enhances outgrowth. Endogenous ERK activity decreases as the EC–tubule distance reaches its plateau, suggesting that the motile program is downregulated as endothelial fronts enter the maintained-separation state. This temporal association does not establish ERK itself as a sensor of EC–tubule distance. Both ERK activity and endothelial motion could instead be controlled by an upstream tubule-associated signal. Thus, ERK is best positioned here as a regulator or readout of the transition between extending and maintained endothelial states. Identifying the upstream cue that links the local tissue environment to ERK dynamics will be important for resolving the molecular basis of the stopping behavior.

A key feature of the follow-and-stop rule is its simplicity. Because the experimentally observed behavior can be expressed using only local motion, directional exploration, and proximity-dependent stopping, it becomes mathematically tractable without requiring detailed assumptions about molecular sensing. The model therefore serves not as a detailed reconstruction of angiogenesis, but as a test of whether the observed local behavior is sufficient to generate larger-scale spatial consequences. Previous modeling of corneal neovascularization has likewise shown that tissue-domain geometry can alter vascular patterning and anastomosis^41^; here, however, the relevant geometry is itself continuously generated by organ morphogenesis. In this reduced description, endothelial connectivity can arise through two complementary routes: stochastic changes in migration direction allow endothelial fronts to explore alternative trajectories, whereas three-dimensional epithelial folding changes the geometry itself and brings otherwise separated fronts into proximity. The model further predicts a trade-off between these mechanisms, such that increasing tissue folding reduces the degree of stochastic exploration required for endothelial encounters. From an engineering perspective, this resembles a system in which relatively simple local control rules operate within a boundary geometry that is continuously being reconfigured.

Our findings therefore suggest a broader way of considering vascularization during organ morphogenesis: endothelial cells need not encode the global architecture of a developing organ if local behaviors are continuously referenced to a morphogenetic tissue structure. Protrusive exploration, approach, and stopping can maintain vascular growth relative to the changing epithelium, while three-dimensional folding alters the spatial opportunities for otherwise distant endothelial fronts to encounter one another. Consistent with this prediction, culture conditions producing greater out-of-plane epithelial folding also showed a higher observed frequency of endothelial connections. However, interface and submerged culture differ not only in epithelial geometry but also in environmental parameters such as oxygen and solute transport and the mechanical constraints imposed by the air–liquid interface. These experiments therefore do not isolate tissue geometry as the sole determinant of endothelial connectivity, and the observed difference should be regarded as directionally consistent with, rather than conclusive evidence for, the model prediction. More generally, the emerging vascular pattern can be viewed as the product of an interaction between molecular regulation of cell motility, stochastic local behavior, and a tissue geometry that evolves over time. This provides a tractable bridge between angiogenic signaling at the cellular scale and organ morphogenesis at the tissue scale, and suggests that changing tissue geometry itself can be an important component of the information landscape within which vascular networks develop.

## MATERIALS AND METHODS

### Animals

Hoxb7-myr-Venus transgenic mice^33^ were used for simultaneous visualization of cell membranes in the epididymal epithelial tubule and developing vasculature. Transgenic mice expressing the hyBRET-ERK-NES biosensor^39^ were used for live imaging of ERK activity. ICR mice were obtained from Japan SLC, Inc. Embryonic stage was defined with the midnight preceding detection of a vaginal plug designated embryonic day 0.0 (E0.0). Epididymides were isolated at E15.5–E17.5 as indicated for individual experiments. All animal experiments were approved by the local institutional animal ethics committees (R22-0765, BR23-1140, and MedKyo 21043) and were conducted in accordance with institutional and national guidelines for the care and use of laboratory animals.

### Antibodies, fluorescent reagents, and pharmacological agents

The following primary antibodies were used: rabbit anti-PAX2 (Invitrogen, #71-6000), rabbit anti-VEGF-A (Abcam, #ab46154), rat anti-CD31 (BioLegend, #102501), rat anti-phospho-histone H3 (Abcam, #ab10543), goat anti-DLL4 (R&D Systems, #AF1389), and goat anti-VEGFR2 (R&D Systems, #AF644). Secondary antibodies included donkey anti-goat Alexa Fluor 555 (Abcam, #ab150134), goat anti-rabbit Alexa Fluor 546 (Thermo Fisher Scientific, #A-11035), and goat anti-rat Alexa Fluor 647 (Thermo Fisher Scientific, #A-21247). Primary and secondary antibodies were used at dilutions of 1:200 and 1:1,000, respectively. Hoechst 33342 (Thermo Fisher Scientific, #62249) was used at a dilution of 1:1,000. Pharmacological agents and recombinant proteins included dimethyl sulfoxide (Nacalai Tescue, #13445-74), PD0325901 (100 nM; FUJIFILM Wako Pure Chemical Corporation, #162-25291), sunitinib malate (5 µM; Tocris, #3768), hEGF (10 ng/mL; Sigma-Aldrich, #E9644), and VEGF-A (100 ng/mL; Merck, #V4512).

### Whole-mount immunofluorescence

Dissected embryonic epididymides were fixed in 4% paraformaldehyde (PFA) in phosphate-buffered saline (PBS) for 2 h at 4°C. After washing with PBS, samples were blocked and permeabilized for 3 h at 37°C in PBS containing 0.1% Triton X-100 and 10% normal goat serum (Abcam, #ab156046) or normal donkey serum (Abcam, #ab166643), depending on the secondary antibodies used. Samples were incubated with primary antibodies overnight at 4°C, washed with PBS, and subsequently incubated with fluorophore-conjugated secondary antibodies overnight at 4°C. Nuclear counterstaining with Hoechst 33342 was performed during the final staining and washing steps. Antibody and Hoechst 33342 dilutions are specified in the preceding section.

### Optical clearing and volumetric fluorescence imaging

For deep three-dimensional imaging, stained or reporter-expressing epididymides were immobilized in approximately 10 µL of 1% agarose on a glass-bottom dish (Greiner Bio-One, #627871) and optically cleared using either benzyl alcohol/benzyl benzoate (BABB; 1:2) or CUBIC2 according to established procedures^42–44^. Volumetric fluorescence images were acquired using a Leica TCS SP8 confocal microscope equipped with HyD detectors and HC PL APO CS2 20× (NA 0.75, WD 680 µm), 40× (NA 1.3, WD 240 µm), or 60× (NA 1.4, WD 140 µm) objectives.

### Ex vivo epididymis culture

Embryonic epididymides were cultured in Dulbecco’s modified Eagle’s medium (DMEM; Nacalai Tesque, #08489-45) or FluoroBrite DMEM (Thermo Fisher Scientific, #A1896701), supplemented with 10% fetal bovine serum (Thermo Fisher Scientific, #12483) and 1% penicillin–streptomycin (Nacalai Tesque, #26253-84). Standard interface culture used a 12-mm Millicell PTFE insert with 0.4-µm pores (Merck Millipore, #PICM01250), with approximately 800 µL of medium placed below the insert. Cultures were maintained at 37°C in a humidified atmosphere containing 5% CO₂.

### Two-photon live imaging

For live imaging, epididymides were cultured in 35-mm glass-bottom dishes (Iwaki, #3910-035) containing approximately 700 µL of culture medium. The supporting legs of the culture insert were immobilized on the dish using approximately 20 µL of 1% agarose to minimize sample drift during imaging. Time-lapse imaging was performed using an incubator-integrated Olympus multiphoton fluorescence microscope equipped with a 25× water-immersion objective (NA 1.05, WD 2 mm; XLPLN25XWMP2, Olympus), with samples maintained at 37°C and 5% CO₂. For imaging of Hoxb7-myr-Venus epididymides, Venus fluorescence was excited at 945 nm using a Mai Tai DeepSee eHP laser (Spectra-Physics). Unless otherwise stated, three-dimensional image stacks were acquired at 2-µm intervals along the z axis and at 5-min temporal intervals. For live ERK activity imaging, epididymides from transgenic mice expressing the hyBRET-ERK reporter were imaged in FluoroBrite DMEM to minimize medium-derived background fluorescence. The reporter was excited at 840 nm. The optical configuration included an RDM690 infrared cut filter and DM505 and DM570 dichroic mirrors (Olympus/Evident); CFP and FRET signals were detected through BA460–500 and BA520–560 emission filters (Olympus/Evident), respectively. Each CFP and FRET image was processed with a 3×3 median filter to suppress shot noise, followed by Gaussian filtering and background subtraction. ERK activity was then calculated pixelwise as the ratio of FRET intensity to CFP intensity. For visualization, the FRET/CFP ratio was represented using a blue–white–magenta lookup table, while image brightness represented FRET-channel fluorescence intensity.

### Local tubule curvature and EC-tubule distance

Three-dimensional PAX2 and CD31 image stacks were converted to 8-bit volumes and analyzed. Individual optical sections were median-filtered, and CD31 images were additionally processed by contrast-limited adaptive histogram equalization. The volumes were smoothed using a three-dimensional Gaussian filter (7×7×3 voxels) and segmented by maximum-entropy thresholding. For epithelial centerline extraction, the largest connected PAX2-positive component was retained, subjected to morphological closing and hole filling, dilated, and skeletonized in three dimensions. For curvature–distance analyses, the traced epithelial centerline was resampled at approximately 100 equally spaced positions, excluding the two terminal points. Local tubule curvature was calculated using a spline curve-based method adapted from previously described approaches^45,46^. Briefly, centerline coordinates were converted to physical spatial units, and consecutive sampling points along the three-dimensional centerline were locally fitted with a cubic spline. Curvature *κ* was calculated from the local derivatives of the fitted spline. Curvature values were smoothed using a seven-point moving average. At each sampled centerline position, EC–tubule distance was defined as the minimum three-dimensional Euclidean distance to the segmented CD31-positive network.

### Geometric analysis of coupled EC-tubule motion

The target epithelial tubule apex and the adjacent endothelial front were tracked through time-lapse image stacks. Tubule-fold displacement, H, was defined as the perpendicular distance from the target tubule apex to the line connecting the two opposing tubule apices on the opposite side of the fold. EC–tubule separation, D, was defined as the Euclidean distance between the tracked EC tip and target tubule apex. The temporal trajectories of H and D were used to describe four operational phases: Ready, during which D remained relatively stable; Go, during which D decreased as the EC front approached the tubule; Stop, corresponding to cessation of the approach; and Keep, during which the reduced EC–tubule separation remained comparatively stable.

### Quantification of endothelial protrusion dynamics and gamma-distribution analysis

Endothelial protrusion dynamics were quantified from Hoxb7-myr-Venus time-lapse images. Individual extension events were defined from the onset of outward protrusion extension until extension ceased and retraction or disappearance began. Extension duration was calculated from the elapsed frames and imaging interval. Each manually measured event contributed a single observation to the analysis. Extension durations were fitted by maximum likelihood to a two-parameter gamma distribution, *p*(*t*) = *t^k^*^−1^exp(−*t*/*θ*)/{Γ(*k*)*θ^k^*}, where *k* and *θ* are the shape and scale parameters, respectively. The fitted parameters were *k*=2.86 and *θ*=43.8 min. To evaluate the distribution of extension durations, gamma, exponential, lognormal, and Weibull models were fitted to the same dataset and compared using the corrected Akaike information criterion. The exponential distribution was included as a reference for a memoryless termination process. Sensitivity analyses were performed using ±15-min bounds around the manually measured durations to account for temporal uncertainty. These adjusted values were not treated as additional independent observations. The estimated gamma shape parameter remained above unity across the tested uncertainty range.

### Quantification of epithelial mitotic density

Mitotic epithelial cells were identified by phospho-histone H3 (pHH3) immunofluorescence using the previously established whole-tissue quantification workflow^42^. PAX2 and pHH3 stacks were median-filtered and smoothed with a Gaussian kernel. The PAX2-positive epithelial compartment was segmented by maximum-entropy thresholding, and the pHH3 channel was segmented by Otsu thresholding. The binary masks were intersected to retain pHH3-positive objects within the PAX2-positive epithelium; connected objects smaller than 10 voxels were removed. The epithelial centerline length was calculated in physical units from the three-dimensional skeleton. Mitotic density was defined as the number of retained pHH3-positive objects divided by centerline length and multiplied by 10, and is therefore reported as mitotic cells per 10 µm of tubule centerline.

### Quantification of three-dimensional tubule shape and endothelial connectivity

Three-dimensional epithelial tubule morphology was quantified from whole-mount PAX2 immunofluorescence images. Image stacks were processed by median and three-dimensional Gaussian filtering, followed by maximum-entropy thresholding, morphological closing, and hole filling. The largest connected epithelial component was retained, dilated, and skeletonized. Following branch removal, the tubule centerline was traced and sampled at intervals of two skeleton voxels. The original image dimensions were 0.7568 µm per pixel laterally and 2 µm per optical section. Local tubule bending was quantified from successive three-dimensional direction vectors calculated over intervals of 20 sampled centerline points. The magnitude of the difference between adjacent unit direction vectors was used as a discrete curvature score, smoothed using a five-point moving average, and averaged along the centerline for each epididymis. To quantify out-of-plane deformation, a least-squares best-fit plane was obtained from the sampled centerline using principal component analysis. The absolute perpendicular distance of each centerline point from this plane was calculated, and the mean distance was used as the out-of-plane deformation measure for each epididymis. Endothelial connectivity was assessed by visual inspection of CD31-positive vascular structures surrounding the epithelial tubule. Epididymides were classified according to the presence or absence of connections between opposing endothelial structures.

### Computational modelling

We developed a minimal three-dimensional model to examine how EC fronts maintain their spatial relationship with a morphogenetic epithelial tubule and how tubule folding promotes connections between opposing endothelial fronts. The epithelial tubule was represented as a cylindrical surface of radius 25 µm surrounding a prescribed, time-dependent centerline. EC fronts were represented as point-like motile agents initially distributed on opposite sides of the straight tubule, with their initial migration directions oriented toward the tubule.

EC movement was governed by a local follow-and-stop rule. Each EC front sampled the epithelial tubule within a forward sensing cone defined by its migration direction and half-angle α. The front advanced at a constant intrinsic speed, v_tip_, when no portion of the tubule was detected within the cone or when the detected tubule surface was farther than the stopping distance, D_stop_=30 µm. Otherwise, forward movement ceased. A minimum surface clearance of 10 µm was imposed to prevent EC fronts from penetrating the epithelial tubule. Thus, proximity to the tubule controlled EC advancement without prescribing a global chemotactic gradient.

For stochastic simulations, directional changes were introduced at renewal intervals drawn independently from the gamma distribution with shape *k*=2.86 and scale *θ*=43.8 min, based on the experimentally measured endothelial protrusion-extension durations. At each renewal, the migration direction was rotated by a normally distributed angular increment with zero mean and standard deviation σ, around a randomly selected azimuth. This parameter therefore represents EC directional fluctuations rather than changes in intrinsic migration speed.

### Parameter estimation for endothelial approach

Planar tubule bending was simulated over 12 h using a prescribed sinusoidal deformation with a maximum amplitude of 120 µm and wavelength of 300 µm. Ten EC fronts were initialized on each side of the tubule. To isolate the effects of intrinsic EC extension speed v_tip_ and sensing-cone half-angle α, directional fluctuations were disabled σ=0. We varied v_tip_ and α, using a simulation time step of 5 min. The mean bilateral EC–tubule separation, *D*, was calculated from the leading EC front and the corresponding epithelial cleft on each side. Late separation was averaged over the final 3 h. Experimental approach speed was estimated by linear regression of individual *D* trajectories during the ‘Go’ phase (−300 to −45 min relative to the onset of the ‘Stop’ phase). Simulated approach speed was calculated from the longest continuous interval in which both leading EC fronts were advancing and *D* was within the experimental ‘Go’-phase distance range. Parameter combinations were evaluated using a normalized mismatch incorporating approach-speed error and a one-sided penalty for late separation exceeding 42 µm. The distance penalty was normalized by 5 µm, and the speed error by the experimental standard deviation. Parameter selection was restricted to combinations with valid bilateral tracking for at least 80% of the final 3 h.

### Three-dimensional folding and endothelial connection in simulations

Tubule morphogenesis was prescribed over 24 h in two successive phases. During the first 12 h, the amplitude of in-plane bending increased smoothly from zero to 120 µm. During the following 12 h, this amplitude was maintained while an orthogonal folding mode developed, with its final relative amplitude determined by sinΦ. Thus, Φ=0 produced purely planar deformation, whereas positive values introduced out-of-plane folding. We fixed v_tip_=0.24 µm/min and α=10, and initialized 20 EC fronts on each side of the tubule. Stochastic directional changes occurred at renewal intervals drawn from a gamma distribution (shape *k*=2.86, scale *θ*=43.8 min). At each renewal, the migration direction was rotated by an angular increment drawn from a zero-mean normal distribution with standard deviation σ, around a uniformly sampled azimuth. A connection was registered when the minimum separation between two opposing EC trajectories within a time step was at most 20 µm, provided that the connecting path did not intersect the epithelial tubule, including a 2-µm clearance. Connected EC fronts were immobilized at their encounter midpoint. We independently varied Φ and σ, and at each parameter combination, 100 stochastic realizations were performed with a 5-min time step. Connection probability was defined as the fraction of simulations containing at least one connection by 24 h. All simulations were implemented in MATLAB (MathWorks).

### Statistical analysis

The number of individual measurements or events is denoted by n, whereas the number of independent embryos or biological replicates is denoted by N. No statistical method was used to predetermine sample size, and no randomization was performed. Exact sample sizes and statistical tests are specified in the corresponding figure legends. We considered p<0.05 statistically significant for two-tailed tests, unless otherwise stated. Significance levels are denoted as follows: * (p<0.05), ** (p<0.01), *** (p<0.001), and n.s. (not significant; p≥0.05).

### Use of artificial intelligence tools

ChatGPT (OpenAI) was used during manuscript preparation to assist with language editing, text restructuring, and debugging of computational code. Explore Science was used as an AI-assisted pre-submission review tool to identify potential issues, reporting, and manuscript positioning. All AI-assisted outputs, code, analyses, and suggested revisions were critically reviewed, tested where applicable, and revised by the authors. The authors take full responsibility for the scientific content and conclusions of the manuscript.

## ACKNOWLEDGEMENTS

This work was supported by the Singapore Ministry of Education Academic Research Fund Tier 2 (MOE-T2EP30223-0010), the National Research Foundation Singapore Mid-sized Grant (NRF-MSG-2023-0001), the Mechanobiology Institute at the National University of Singapore through the Research Centre of Excellence programme funded by the National Research Foundation and Ministry of Education, Singapore, the Kao Foundation for Arts and Sciences, the Takeda Science Foundation, and the Hakubi Fellowship of Kyoto University. We thank Taiji Adachi and Michiyuki Matsuda for providing access to laboratory resources and our laboratory members for their valuable comments on the manuscript.

## CODE AVAILABILITY

Relevant code for this study will be available at a GitHub repository.

## VIDEO LEGENDS

**Video 1.** Three-dimensional organization of the developing vasculature during epididymal tubule morphogenesis. Three-dimensional visualizations of whole-mount immunofluorescence images of mouse epididymides at E17.5. CD31-positive endothelial cells (magenta) and the PAX2-positive epithelial tubule (blue) are shown, illustrating the progressive extension of the vascular network into the organ interior as the epithelial tubule folds. Related to Figure 1A.

**Video 2.** Three-dimensional reconstruction of the endothelial network and epithelial tubule. Representative three-dimensional segmentation of the CD31-positive endothelial network and PAX2-positive epididymal tubule from whole-mount fluorescence images. The reconstruction illustrates their spatial organization and the epithelial centerline used to quantify local tubule bending and EC–tubule distance. Related to Figure 1B.

**Video 3.** Dynamic extension and retraction of endothelial protrusions toward the epithelial tubule. Ex vivo two-photon time-lapse imaging of a Hoxb7-myr-Venus epididymis showing repeated extension and retraction of filopodia-like protrusions from an endothelial front toward the adjacent epithelial tubule. The sequence illustrates the dynamic exploration of the local epithelial environment by endothelial protrusions. Related to Figure 3D,E.

**Video 4.** Endothelial fronts approach and maintain their separation from a folding epithelial tubule. Long-term ex vivo two-photon imaging of an E15.5 epididymis showing endothelial extension alongside progressive epithelial tubule folding. Following an initial delay, the endothelial front advances toward the developing fold and subsequently maintains a relatively stable separation from the tubule despite continued epithelial deformation. Related to Figure 4A–C.

**Video 5.** Endothelial retraction accompanies a local change in epithelial tubule geometry. Representative ex vivo time-lapse imaging showing a local change in epithelial folding accompanied by retraction of the neighboring endothelial front. The sequence illustrates the dynamic repositioning of endothelial structures as the epithelial geometry changes. Related to Figure 4D.

**Video 6. Three-dimensional visualization of ERK activity in the developing epididymis.** Three-dimensional visualization of the hyBRET-ERK reporter signal in an embryonic epididymis. ERK activity is represented by the FRET/CFP ratio using a blue–white–magenta color scale, illustrating the spatial distribution of ERK signaling within the developing tissue. Related to Figure 5.

**Video 7.** Live imaging of endothelial ERK activity during approach toward the epithelial tubule. Ex vivo two-photon time-lapse imaging of ERK activity in a hyBRET-ERK reporter epididymis. The sequence illustrates spatially heterogeneous endothelial ERK signaling and its temporal decline as the endothelial front transitions from active approach to maintenance of its separation from the epithelial tubule. Related to Figure 5F–H.

**Video 8.** Computational simulation of endothelial extension and connection during epithelial tubule folding. Three-dimensional visualization of the minimal follow-and-stop model, in which endothelial fronts extend toward a deforming epithelial tubule and cease forward advancement upon reaching a prescribed proximity range. The simulation illustrates how epithelial folding and stochastic changes in endothelial migration direction influence encounters and connections between opposing endothelial fronts. Related to Figure 6.

**Video 9.** Three-dimensional comparison of epithelial folding and vascular organization under interface and submerged culture conditions. Side-by-side, rotating three-dimensional reconstructions of embryonic epididymides following 24 h of ex vivo interface culture (left) or submerged culture (right). The PAX2-positive epithelial tubule is shown in blue and the CD31-positive endothelial network in magenta. Rotation reveals the predominantly planar tubule configuration under interface culture and the more pronounced out-of-plane folding under submerged culture, together with their associated vascular architectures. Related to Figure 7.

