## Supplementary note for "Geometric coordination of endothelial invasion and vascular patterning during epithelial morphogenesis"

### 1 Reduced geometric–stochastic encounter model

The main text proposes that out-of-plane epithelial folding and stochastic changes in endothelial migration direction provide two routes to connection. Here we formalize this idea with a minimal encounter model and derive the contour relation quoted in the main text. The purpose of the reduced model is not to reproduce the complete tip trajectories, but to isolate the balance between *geometric access* and *stochastic exploration* near the connection boundary.

Consider a prospective pair of endothelial tips. Let $b(\Phi) \geq 0$ denote their effective deterministic transverse miss distance and $a > 0$ an effective capture half-width. We assume that, close to the connection transition, out-of-plane deformation reduces the geometric gap according to

$$b(\Phi) \simeq b_0 - \Lambda \sin\Phi, \qquad \Lambda > 0, \tag{1.1}$$

where $b_0$ is the miss distance without out-of-plane deformation and $\Lambda$ measures the geometric contribution of folding. This is a local approximation for an accessible encounter route; it does not require every Euclidean tip–tip distance to decrease monotonically with $\Phi$. In the absence of directional fluctuations, connection becomes geometrically possible when $b(\Phi) \leq a$, defining

$$\sin\Phi_c = \frac{b_0 - a}{\Lambda}. \tag{1.2}$$

Thus $\Phi_c$ is the deterministic geometric connection threshold of the reduced model.

Directional fluctuations broaden the region explored by the tips. Suppose a tip moves with speed $v_{\mathrm{tip}}$ and its direction is renewed with mean interval $\tau = k\theta$, where $k$ and $\theta$ are the shape and scale of the renewal-time distribution. For small angular changes with standard deviation $\sigma$, the relative transverse displacement of two independently fluctuating tips over an effective exploration time $T_{\mathrm{eff}}$ has the leading-order variance

$$s^2 \simeq \frac{v_{\mathrm{tip}}^2 \sigma^2 T_{\mathrm{eff}}^3}{3k\theta}, \qquad s = K\sigma, \qquad K = \frac{v_{\mathrm{tip}} T_{\mathrm{eff}}^{3/2}}{\sqrt{3k\theta}}. \tag{1.3}$$

Here $s$ is the characteristic transverse exploration width. This scaling follows from integrated angular diffusion and is intended for the small-turn, many-renewal regime.

To translate this exploration width into a connection probability, we write the effective transverse offset at the encounter as

$$X = b(\Phi) + Y, \qquad Y \sim \mathcal{N}(0, s^2), \tag{1.4}$$

and take connection to occur when $|X| \leq a$. The corresponding single-route capture probability is

$$q(\Phi, \sigma) = \mathcal{F}\left(\frac{a - b(\Phi)}{s}\right) - \mathcal{F}\left(\frac{-a - b(\Phi)}{s}\right), \tag{1.5}$$

where $\mathcal{F}$ is the standard normal cumulative distribution function. Close to the low-fluctuation accessibility boundary, where $b(\Phi) > a$ and the far edge of the capture interval contributes little, this reduces to

$$q \simeq \mathcal{F}\left(\frac{\Lambda \sin\Phi - (b_0 - a)}{K\sigma}\right). \tag{1.6}$$

For any fixed connection-probability contour, the corresponding value of the Gaussian argument is constant. Denoting its positive magnitude by $c_p$ gives the central geometry–fluctuation balance

$$\boxed{\underbrace{\Lambda \sin\Phi}_{\text{geometry}} + \underbrace{c_p \frac{v_{\text{tip}} T_{\text{eff}}^{3/2}}{\sqrt{3k\theta}}\,\sigma}_{\text{stochastic exploration}} \gtrsim \underbrace{b_0 - a}_{\text{gap to overcome}}} \tag{1.7}$$

Here $c_p$ collects the probability threshold into an order-one constant; the detailed mapping between a local capture probability and the system-level probability $p$ is not needed for the normalized contour derived below because $c_p$ cancels. Equation (1.7) makes the physical interpretation explicit: below the deterministic connection threshold, folding and directional fluctuations compensate for one another. Increasing $\Phi$ reduces the stochastic exploration needed to bridge the remaining gap.

Setting the balance to equality gives the fluctuation amplitude required to remain on a fixed probability contour,

$$\sigma_p(\Phi) \simeq \frac{b_0 - a - \Lambda \sin\Phi}{c_p K}. \tag{1.8}$$

At $\Phi = 0$,

$$\sigma_p(0) = \frac{b_0 - a}{c_p K}. \tag{1.9}$$

Using Eq. (1.2) to eliminate the unknown geometric scale $b_0 - a = \Lambda \sin\Phi_c$, division of Eq. (1.8) by Eq. (1.9) yields

$$\boxed{\frac{\sigma_p(\Phi)}{\sigma_p(0)} \simeq 1 - \frac{\sin\Phi}{\sin\Phi_c}} \qquad (0 \le \Phi \lesssim \Phi_c). \tag{1.10}$$

This is the analytical relation used in the main text. Its normalized form is independent of $v_{\text{tip}}$, $T_{\text{eff}}$, $k$, $\theta$, and the probability-dependent coefficient $c_p$; these quantities determine the absolute fluctuation scale $\sigma_p(0)$ but cancel from the contour shape. The prediction is therefore that the low-fluctuation transition boundary is approximately linear when plotted against $\sin\Phi$ and terminates as the deterministic geometric threshold $\Phi_c$ is approached.

The same reduced model also clarifies why the compensation cannot continue indefinitely. If the deterministic trajectory is already inside the capture region, $b(\Phi) < a$, additional directional dispersion broadens the trajectory away from an already accessible target and can reduce rather than increase connection probability. Thus geometry and stochasticity are complementary below the accessibility threshold but can become antagonistic once geometric access is sufficient. Equation (1.10) is consequently intended only for the descending, low-fluctuation transition boundary, not for the entire parameter plane.

### 2 Comparison with the full three-dimensional simulations

We next asked whether the simple analytical prediction in Eq. (1.10) captures the connection boundary generated by the full endothelial-tip simulations described in the main text. Unlike the reduced encounter model, the simulations explicitly include the deforming epithelial tube, forward-cone sensing and GO/STOP motion, stochastic three-dimensional direction renewal, geometric obstruction by the tube, and path-dependent tip–tip connection. They therefore provide an independent numerical test of whether the predicted geometry–fluctuation balance survives these additional constraints.

Because out-of-plane folding is introduced after the initial planar-folding stage, connections can in principle arise before or after the out-of-plane deformation develops. The analytical model does not attempt to reproduce this time dependence. Instead, we compare only the *shape of the final connection-probability boundary* in the $(\Phi, \sigma)$ plane.

In the simulated parameter map, zero-noise trajectories remain unconnected through the sampled $\Phi = 15.75°$ and connect at the next sampled angle, 18°, placing the deterministic transition within this interval. We then extracted the first upward crossing of selected connection-probability levels as $\sigma$ increased and fitted the low-fluctuation branch with

$$\sigma_p(\Phi) = A_p \left(1 - \frac{\sin\Phi}{\sin\Phi_c}\right), \tag{2.1}$$

which is the dimensional form of Eq. (1.10) with $A_p = \sigma_p(0)$.

**Table S1:** Descriptive fits of the analytical contour shape to the simulated low-fluctuation connection boundaries. Angles and root-mean-square errors (RMSE) are in degrees. The fitted parameters are estimated from the numerical map and are used only to assess the predicted functional form.

| $p$ | $A_p$ | fitted $\Phi_c$ | RMSE | $R^2$ |
|---|---|---|---|---|
| 0.25 | 13.48 | 18.25 | 0.76 | 0.970 |
| 0.50 | 20.14 | 18.31 | 0.93 | 0.979 |
| 0.75 | 29.48 | 19.88 | 3.53 | 0.857 |

The $p = 0.25$ and $p = 0.50$ contours closely follow the predicted approximately linear dependence on $\sin\Phi$, with fitted $\Phi_c$ values near the deterministic transition identified from the parameter grid. The agreement is weaker for $p = 0.75$, consistent with the reduced model being most appropriate near the low-fluctuation accessibility boundary rather than at high connection probability. Fixing $\Phi_c = 18°$ and fitting only the amplitude gives $R^2 = 0.969$ for $p = 0.25$, 0.978 for $p = 0.50$, and 0.833 for $p = 0.75$.

Together, the analytical model and simulations support a simple interpretation of the parameter map: out-of-plane epithelial folding reduces the geometric gap between otherwise disconnected endothelial trajectories, whereas stochastic directional fluctuations broaden the space explored by the tips. Near the transition, these two effects compensate according to Eq. (1.10); once geometric access becomes sufficient, additional fluctuations need not remain beneficial. The analytical calculation therefore provides a minimal geometric explanation for the connection boundary observed in the full simulations without attempting to reproduce the complete path-dependent dynamics.

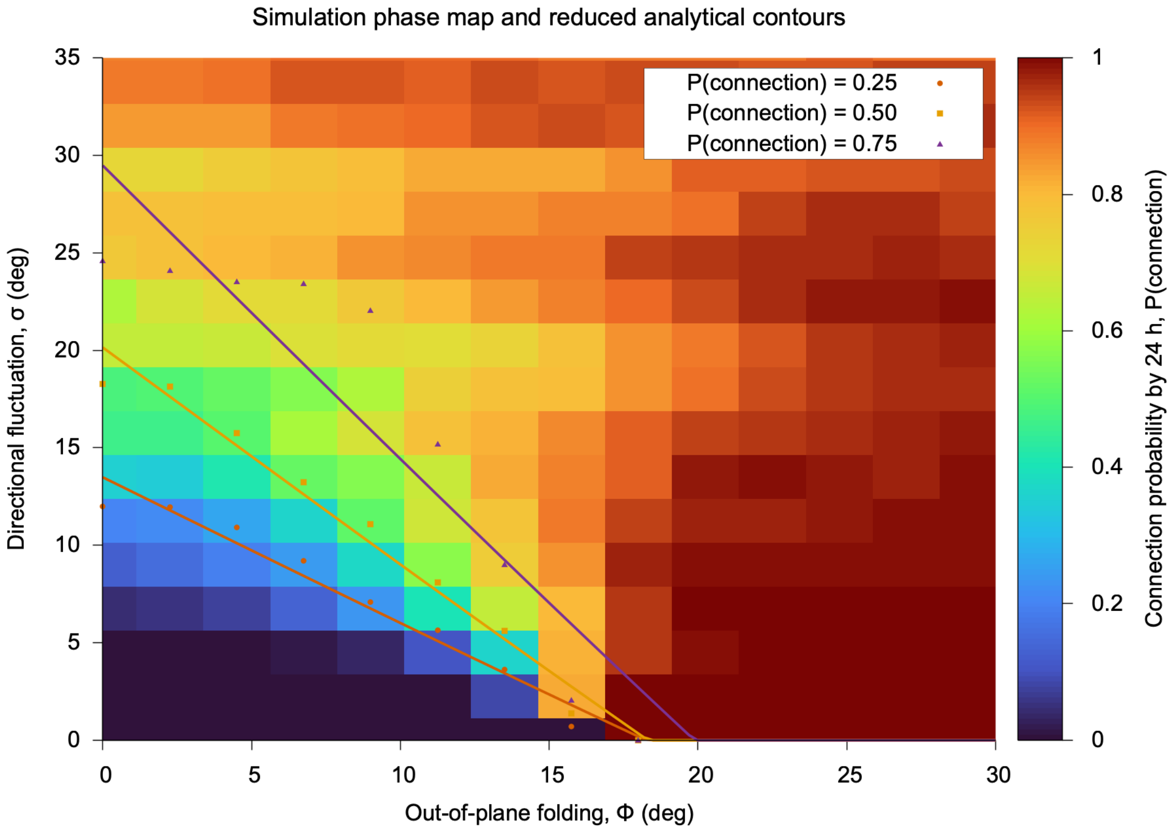


**Figure S1:** Connection probability in the full three-dimensional simulations (background) compared with the first upward probability crossings (open symbols) and descriptive fits of Eq. (2.1) to the low-fluctuation branches (dashed curves). The analytical form captures the descending $p = 0.25$ and $p = 0.50$ boundaries more closely than the higher-probability $p = 0.75$ boundary.
